# Characterization of *B646L* (p72) gene and the resistance pattern of African swine fever Virus tolerant to indigenous Doom Pig breeds of India

**DOI:** 10.1101/2023.05.25.542278

**Authors:** Pranab Jyoti Das, Joyshikh Sonowal, Gyanendra Singh Sengar, Seema Rani Pegu, Rajib Deb, Satish Kumar, Santanu Banik, Swaraj Rajkhowa, Vivek Kumar Gupta

## Abstract

African Swine Fever (ASF) has affected all pig breeds in North-East India since 2020, except Doom pigs, a unique indigenous breed from Assam and the closest progenitor to Indian wild pigs, resulting in significant economic losses for pig farmers in the region. Based on the complete sequences of the *B646L* (p72) gene, it has been determined that the virus responsible for the outbreak is ASFV genotype II. The further characterization of three complete sequences of the *B646L* (p72) gene established 100% identical with other existing sequences of different parts of the world as well as confirmed that there is no co-circulation of different genotypes of ASFV in India except genotype II. Present studies also corroborate that *MYD88, LDHB* and *IFIT1* were important genes of the immune system involved in the pathogenesis of ASFV. The differential expression patterns of these genes in ASFV-infected survived, and healthy Doom breed pigs, compared to healthy control pigs, were studied to distinguish the expression pattern at different stages. The hardiness and ability of the Doom pig to withstand common pig diseases, along with its genetic resemblance to wild pigs, make it an ideal candidate for studying tolerance to ASFV infection. So, the present study investigates the natural resistance to ASF in Doom pigs from an endemic area in North-East India to support the proposition that Doom pigs can co-exist with virulent ASFVs recently break in North-East India. The results of this study also provide important molecular insights into the regulation of the ASFV-tolerant gene.

**Importance:** Studying the natural resistance to African Swine Fever (ASF) in Doom pigs from North-East India holds crucial importance. ASF has inflicted significant economic losses on pig farmers in the region, necessitating the identification and comprehension of factors contributing to resistance and tolerance in specific pig breeds such as Doom pigs. Understanding the molecular mechanisms and genetic factors associated with ASFV tolerance could help in breeding programs and the selection of resilient pig breeds, ultimately aiding in disease control efforts.

## Introduction

African Swine Fever (ASF) virus (ASFV), the causative agent of ASF is a large, enveloped, icosahedral, double-stranded DNA virus which belongs to the genus *Asfvirus*, family *Asfaviridae* and order *Asfuvirales* (1). Current investigation indicates that capsid protein P72 of ASFV is very important for the diagnosis and future vaccine development (2). As the ASFV continues to spread to pig populations globally different protection and antiviral agents are being developed to combat the virus. Previously, researchers also showed that warthog which is a wild member of the pig family found in grassland, savannah, and woodland in sub-Saharan Africa carries certain transcriptomic factors that show resistance to ASFV (3). Another recent study shows that Kenya domestic pigs which are pure indigenous pigs of Kenya also show the same kind of resistance to ASFV (4, 5). The detection of ASFV was also seen to correlate with pig genotype, with a higher proportion of pigs with low local African ancestry testing positive for the virus. A comparative study on selection signatures between the wild African and local domestic African pigs indicates the absence of broad patterns of selection that are similar between the domestic and wild pigs (4). But this may be due to the small number of markers that are amplified for warthog and bush pig, limiting the ability to detect selection signatures with good accuracy. But revealing some evidence of introgression of wild pigs into domestic Kenya pigs indicates that a natural crossing over was taken place at some point in time. Since it is already established that wild African pigs are resistant to ASFV, the presence of viable hybrids between domestic and wild pigs or identifying close progenitors of the wild pigs presents an opportunity to further characterize the genetic basis of ASFV tolerance or resistance. These indigenous pigs particularly found in Africa and Asia are the natural habitat of pigs’ domestication and origination and they are more closely related to their wild ancestors than the modern European and American breeds. Few studies also show that African pigs have a composite ancestral relation with haplotypes from Asian counterparts.

In the Asian context, particularly, in the Northeastern region of India which is one of the main hotspots for pig origination and domestication has some prized indigenous pig breeds. Among total registered pig breeds (*viz*. Ghungroo, Niang Megha, Tanye Vo, Zovawk, Doom, and Mali) from NE India, Doom pigs are very much prevalent in Assam and are the closest progenitor of Indian wild pigs (unpublished data). Moreover, this pig has been used in the maternal line for the production of the HDK75 crossbred variety of pigs. Along with other crossbred varieties like Rani, Asha, and Lumsniang; the HDK75 crossbred variety is very popular among pig farmers not only for their production and reproduction performance but also due to hardiness and coping power with common pig diseases.

The recent outbreak of ASFV in the region causes huge economic losses to pig farmers in the region and all kinds of pigs of irrespective exotic, crossbred and indigenous pigs are affected by this virus except Doom pigs. It is estimated the direct losses due to the loss of animals due to ASFV in India is INR 2.76 billion (US$ 37.32 million) (6). But this estimation was only up to July 2021; the disease was more prominent and havoc in the region in a recent couple of years. But interestingly there is nil report of ASFV infection in Doom pigs as well sporadic report of certain pigs irrespective of any breed survived ASVF infection. So, the present study was undertaken on the Doom breed of pig which is the closest progenitor of wild pigs to analyse the key genetic biomarker/ genes expression pattern that confers tolerance to ASFV in Doom and ASFV survived pigs of India as well as to analyse genetic sequences of the most dominant structural component of the ASFV the major capsid protein p72 within the infected region.

## 2. Materials and methods

### 2.1 Collections of blood and tissue samples

Tissue and blood samples of pigs were collected by the trained person from the Doom, ASFV susceptible and ASFV tolerant pigs (Pig Farm ICAR-National Research Centre on Pig, Rani Guwahati and KVK Goalpara and from the infected zone). Whole blood-EDTA and serum samples were divided into different groups according to infected animals as (i) infected and died and (ii) infected and survived (recovered). Samples were also collected from healthy non-infected animals of (iii) indigenous breed (Doom) and (iv) exotic breed (Yorkshire). The collected blood samples were put into the RNAlater tube and stored at -80 ^0^C for further use.

### 2.2 DNA extraction

Blood samples of all categories of pigs were subjected to extract the genomic DNA using DNeasy Blood & Tissue Kits (QIAGEN, Germany) manufacturer’s protocol. The quality and concentration of isolated DNA samples were checked spectrophotometrically (Eppendorf, Hamburg, Germany). Extracted DNA samples were stored at -20°C till further use.

### 2.3 RNA isolation and cDNA synthesis

Total RNA from resistant, susceptible and normal pigs were isolated from PBMC using RNeasy Plus Universal Mini Kit (QIAGEN, Germany) according to the manufacturer’s instructions. The quality and quantity of the isolated total RNA were measured using a spectrophotometer and synthesised the cDNA using QuantiTect Reverse Transcription Kit (QIAGEN, Germany). Synthesized cDNA was kept at -80°C till further use.

### 2.4 Synthesis of Oligos

In this study, a total of five genes’ oligonucleotides *viz. LDHB* (Lactate Dehydrogenase B), *IFIT1* (Interferon-induced protein with tetratricopeptide repeats 1), *MYD88* (Innate immune signal transduction adaptor), *GAPDH* and *ACTB* were designed using available online software Primer3 plus (https://www.bioinformatics.nl/cgi-bin/primer3plus/primer3plus.cgi) (**Table 1**). Introns spanning primers for *GAPDH* and *ACTB* genes were designed for RT-PCR so that DNA contamination can also be detected.

**Table 1:**
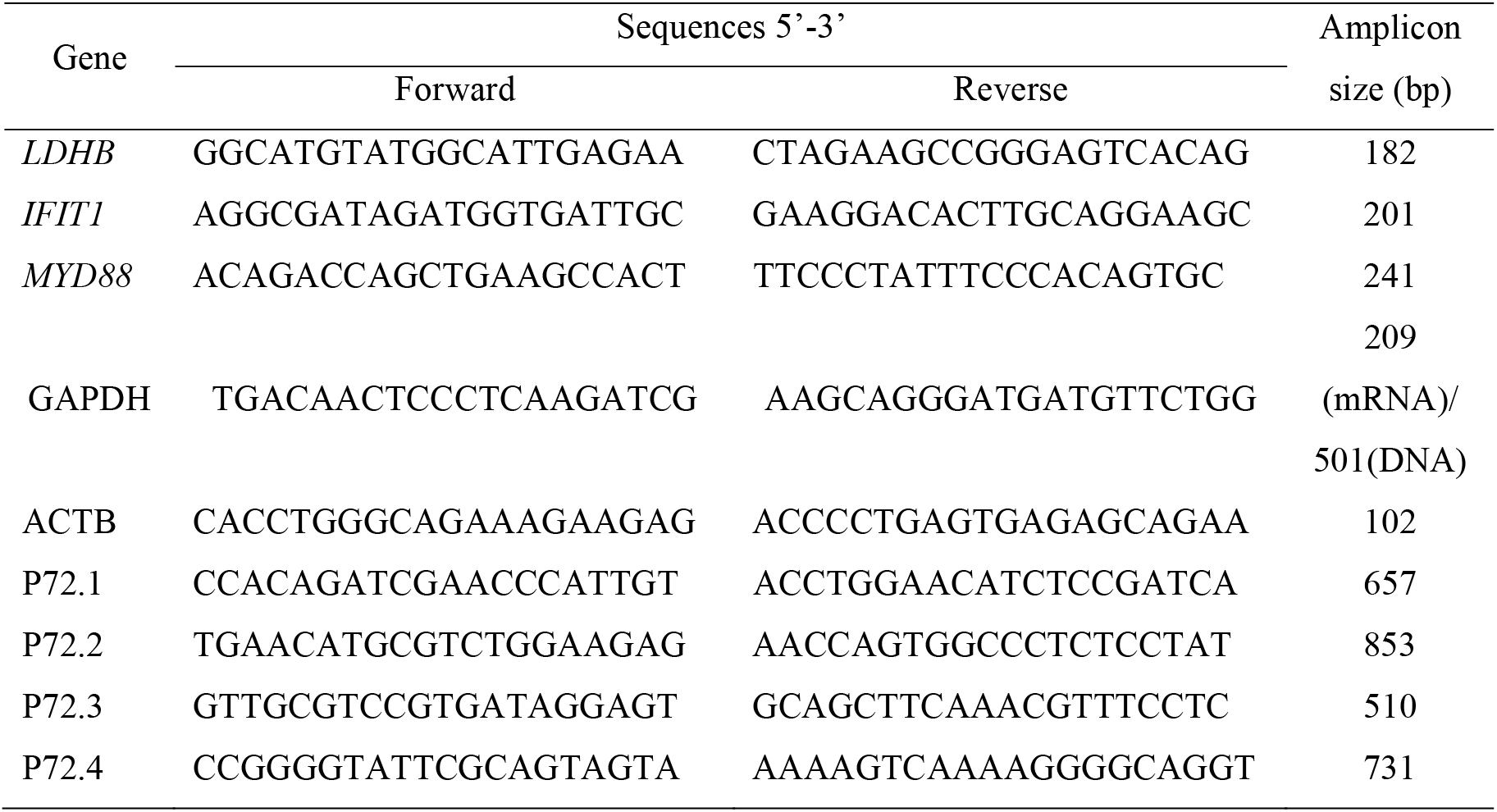
Details of Oligos used in the current study

### 2.5 Confirmation of the ASFV infection

The isolated DNA of each sample was used to confirm the presence of ASFV. For PCR amplification, the primers forward 5′-CCGGGGTATTCGCAGTAGTA -3′ and reverse 5′-GCAGCTTCAAACGTTTCCTC -3′ were used at an annealing temperature of 58°C. Additionally, three other pairs of primers mentioned in Table 1 were also used for the detection of the ASFV p72 gene.

### 2.6 Primer designing, Amplification, Sequencing and phylogenetic analysis of the complete *B646L* (p72) gene of ASFV

A total of four pairs of primers *viz*. P72.1, P72.2, P72.3 and P72.4 (Table.1) were designed covering total 2255bp covering 131bp 5’ upstream, 183bp 3’ downstream and 1941bp *B646L* (p72) gene using referral sequence ID OK236383 and Primer3 Plus software. Amplification of all the fragments size of 657bp, 853bp, 510bp and 731bp covering complete *B646L* (p72) genes was carried out by PCR in a 50 *μ*L of reaction volume containing 1× Dream *Taq* buffer, 5 U DNA polymerase (Invitrogen, Carlsbad, CA, USA), 0.2mM dNTPs (Invitrogen), 25 pmol primers and 100ng DNA. The reactions were carried out as follows: first denatured the DNA was at 94°C for 5 min, repeat a cycle of 94°C for 30 s, 58*/*60°C for 30 s, 72°C for 1 min for 30 cycles and a final extension of 10 min at 72°C in a thermal cycler. This was followed by standard double-stranded sequencing reactions performed using 50ng of purified PCR product, 4pmol of primer and a BigDye Terminator-ready reaction kit (Applied Biosystems, Foster City, USA). Cycle sequencing was carried out using both forward and reverse primer of each fragment in a GeneAmp9600 thermal cycler (Perkin Elmer, Waltham, USA) employing 30 cycles at 96°C for 10 s, 49−50°C for 5 s and 60°C for 4 min. Extended products were purified by alcohol precipitation followed by dissolving in Hi-Di formamide and analysing in an ABI3700 automated DNA Analyzer (Applied Biosystems). Raw sequences are assumable and aligned using NCBI blast and submitted to NCBI databank for accession numbers. Subsequently, another nine *B646L* (P72) sequences with accession numbers OP331311, MN793051, MZ054172, MW296946, MW296945, OL692744, OL692743, LR899193 and MW049116 were downloaded NCBI (http://www.ncbi.nlm.nih.gov/) used for construction phylogenetic tree, distance matrix and alignment report using all three new sequences of present study. The phylogenetic tree, distance matrix and alignment report were performed using MEGA v11 (7). Consensus sequence and gap weight were analysed and Distance metrics between the samples were calculated by using MegAlign of DNAstar (8). Mismatch analysis values were estimated using 100% bootstrap values. These values were used to reconstruct the neighbour-joining tree by comparing different *B646L* (p72) gene sequences available at the NCBI databank.

### 2.7 Persistency of ASFV antibody

Serum samples of infected pigs were collected during infection and recovered pigs on the 0^th^, 14^th^, 28^th^, and 35^th^-day post recovery and the persistency of antibody against ASFV was confirmed by INgezim ASF iELISA kit (INGENSA, Spain) as per manufacturers protocol.

### 2.8 Quantitative analyses of ASFV resistance gene

Quantitative real-time PCR (qPCR) was carried out to validate differentially expressed transcripts of resistant, susceptible and tolerant pigs using an SYBR green in AriaMx Real-Time PCR system (Agilent, US). The qPCR was performed in a 10μL reaction volume containing 1μL cDNA (50ng) as the template, 5μL SYBR Green PCR Master Mix, 1μL each of the forward and reverse primers (2 pmol), and 3μL H_2_O using the following cycling parameters: 40 cycles at 94°C for 5 s and 60°C for 10s. A melting curve was constructed at the end of the PCR for 5 s from 60°C to 95°C to identify unique PCR products amplified during the reaction. The gene *ACTB* was used as a reference gene. The qPCR reaction was repeated three times independently and the data were analyzed by the relative quantification method using 2^-ΔΔCt^. The significance of differences was analyzed using Student’s t-test analysis of variance (ANOVA) with p < 0.001 considered significant.

### 2.9 Ethics statement

Procurement of blood and tissue samples was performed following the approval of the Institute Animal Ethics Committee of the Indian Council of Agricultural Research-National Research Centre on Pig, Rani, Guwahati, Assam, India. The approved animal use protocol number is NRCP/IAEC/1658/2023-24/90 dated 24.04.23.

## 3. Results

### 3.1 Confirmation of ASFV infection in infected and recovered animal

The presence of 156bp size of amplified product {*B646L* (p72) gene} in infected animals confirmed the presence of ASFV in the sample (Fig. 1A). In the survived animal no band was found, which indicates the samples negative for ASFV (Fig. 1B). *GAPDH*, product size 156bp was amplified in all the infected and survived samples (Fig. 1).

**Fig. 1:**
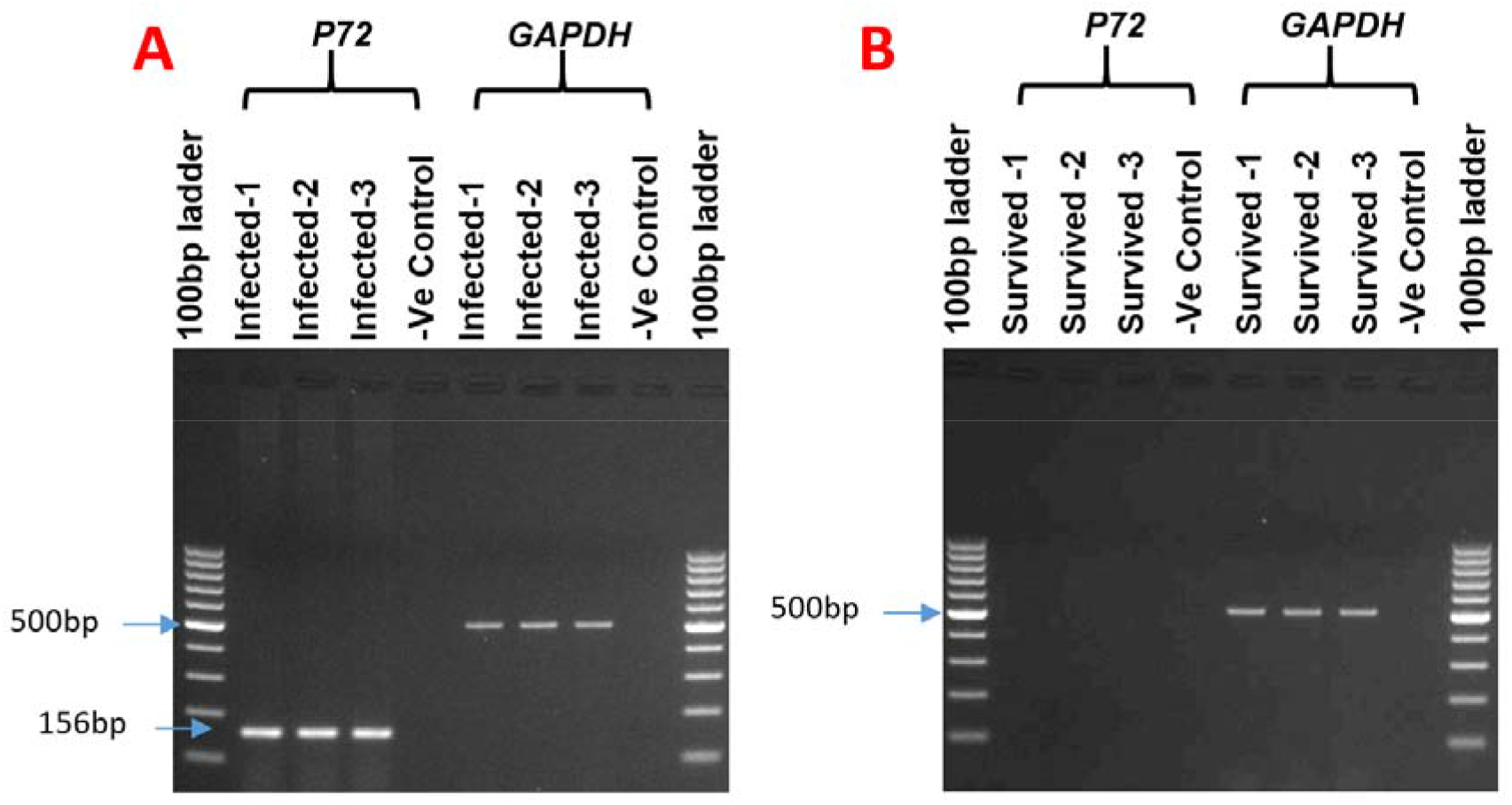
Agarose gel electrophoresis results confirming the PCR amplification of the ASFV p72 gene (156bp) and GAPDH (501bp) from blood samples of pigs that were either infected with ASFV or survived. The infected pig samples (1, 2, 3) exhibited bands at the respective positions indicating the presence of ASFV p72 gene products (A), whereas the survived pig samples (B) did not show any band at that position. Both the infected and healthy pig samples displayed the endogenous control band of GAPDH at the appropriate position.

### 3.2 Characterization of African swine fever Virus *B646L* (p72) Genotype II from Northeast India

Three blood/tissue samples from different locations of ASFV infected area of Northeast India (Fig. 2) were subjected to viral DNA isolation and amplification complete *B646L* (p72) amplification with four sets of primers (Table. 1, Fig. 3). Amplified products were used for sequencing 2255bp of *B646L* (p72) gene along with 131bp 5’upstream, 183bp 3’downstream and 1941 complete *B646L* (p72) gene by standard double-stranded sequencing reaction and analysed in ABI3700 automated DNA Analyzer (ABI). All three *B646L* (p72) gene sequences of ASFV were submitted to the NCBI databank and got the accession numbers ON875961, OP796667 and OP796668.

**Fig. 2:**
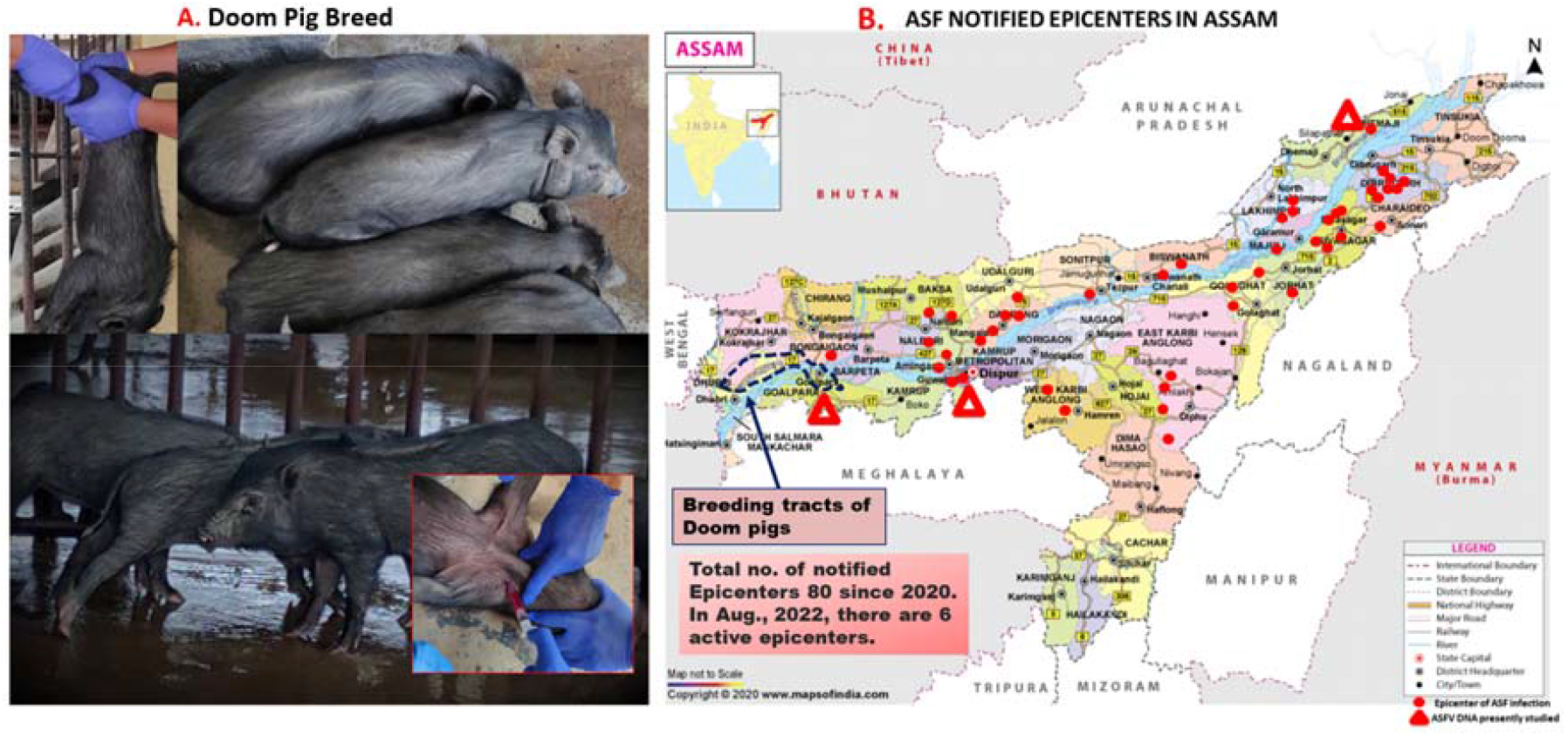
A. Representation of Doom breed of pigs used in the experiment, B. Representation of the geographical area of pig sample collection for the present study. The map displays the breeding tracts of Doom breed pigs and the epicentres of African swine fever from 2020 to 2022 in Assam, India.

**Fig. 3:**
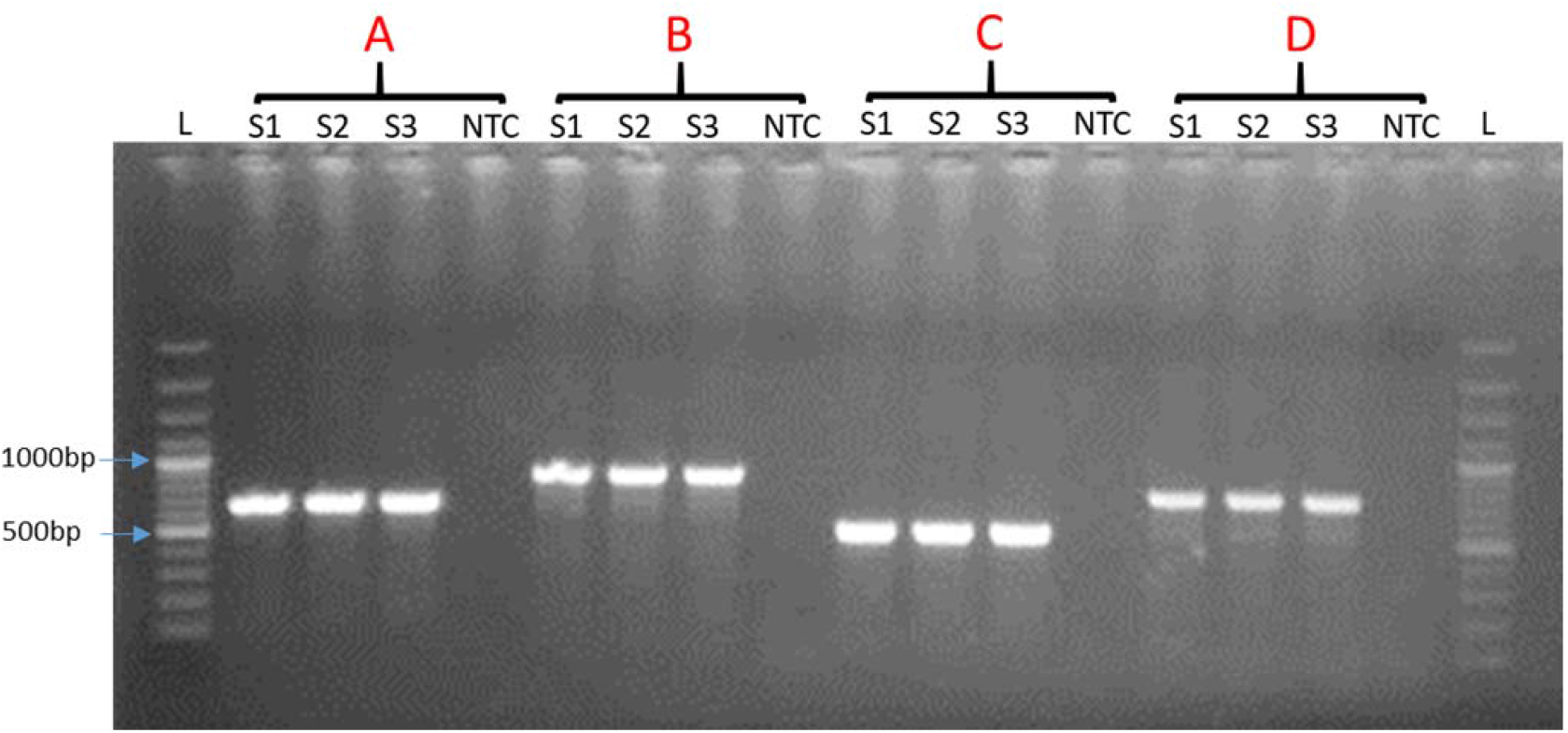
Amplification of different fragments of (A) 657bp, (B) 853bp (C) 510bp and (D) 731bp covering total 2255bp that contains 131bp 5’ upstream, 183bp 3’ downstream and 1941bp *B646L* (p72) gene in S1, S2, S3 sample. Here, L=100bp plus ladder, NTC= Negative control

Subsequently this three these sequences were aligned with available *B646L* (p72) gene sequences Genotype II ASFV of OP331311_China, MN793051_Vietnam, MZ054172_Nigeria, MW296946_Estonia, MW296945_Armenia, OL692744_India, OL692743_India, LR899193_Germany and MW049116_Korea (http://www.ncbi.nlm.nih.gov/). The Phylogenetic analysis, distance matrix and alignment report confirmed that all the *B646L* (p72) sequences from a different location in Northeast India belong to Genotype II ASFV and share 100% sequence similarity with other analysed sequences from China, Vietnam, Nigeria, Estonia, Armenia, India, Germany and Korea (Fig. 4).

**Fig. 4:**
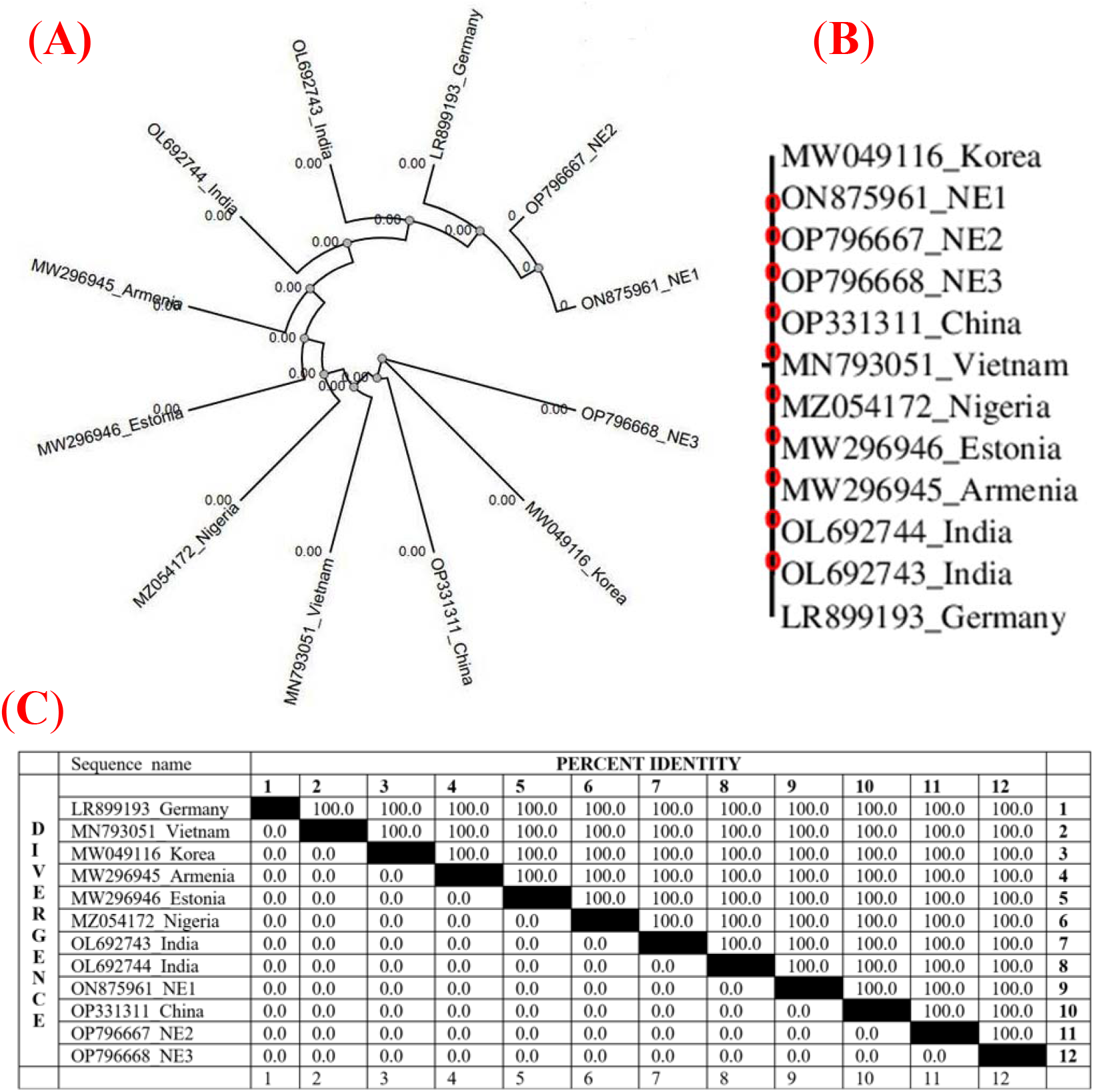
(A) Radial Phylogram and (B) Phylogram with branch length of ON875961_NE1, OP796667_NE2, OP796668_NE3 (Present study), OP331311_China, MN793051_Vietnam, MZ054172_Nigeria, MW296946_Estonia, MW296945_Armenia, OL692744_India, OL692743_India, LR899193_Germany and MW049116_Korea are position in a single cluster with 0.00 branch length belongs to Genotype II ASFV. (C) Pairwise distance matrix calculated ClustalW showing per cent identity and divergence of VP72 gene sequences of ASFV genotype calculated using MegAlign program. All the tested sequences of the present study viz. ON875961_NE1, OP796667_NE2, and OP796668_NE3 show 100% similarity with existing other sequences of VP72 gene sequences.

### 3.3 Presence of antibodies in infected and recovered pigs

When the first clinical symptoms appeared, none of the infected animals had any antibodies, but following severe infection, two of them tested positive for ASFV antibodies, which were followed by the animal’s death (Fig. 5A). The animals tested negative for ASFV in the PCR and lateral flow assay tests on the first post-recovery (PR) day (0th day PR). Additionally, from the first day after recovery forward, positive ASFV antibodies were discovered in the recovered animals, and the persistence of ASFV antibodies was examined up to the 35^th^ day after recovery (Fig. 5B). In this time frame, the antibody’s peak was found on the 14^th^ day of PR. The ELISA report emphasizes that the animals used for the present study recovered from ASFV and have persisted antibodies against ASFV. This means our report validated the targeted gene expression studies in ASFV-resistant animals.

**Fig. 5:**
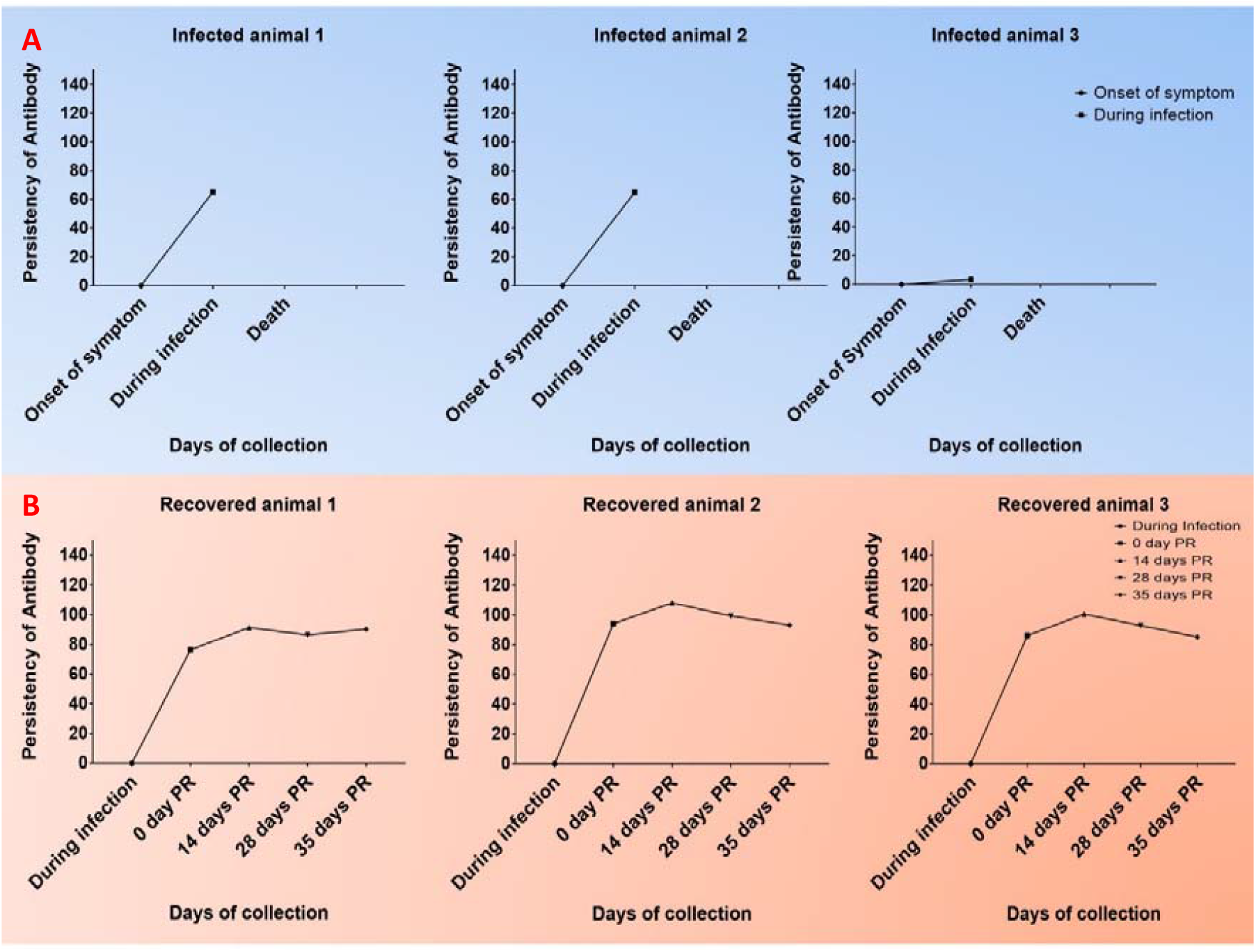
A graphic depiction of the persistence of antibodies against the African swine fever virus (ASFV) in each infected pig (A. infected animals numbered 1, 2, and 3), as well as in animals that have recovered (B. recovered animals numbered 1, 2, and 3).

### 3.4 Quantitative analyses of ASFV resistance gene

According to the qPCR analysis, the relative fold change of *MYD88* expression in ASFV-infected pigs was 1.22 fold, while in survived pigs it was 0.69 fold, and in healthy Doom, it was 0.17 fold compared to healthy control (which was set as 1 fold) (Fig. 6 A2). These results suggest that *MYD88* expression is upregulated during ASFV infection, but is downregulated in pigs that survive the infection. In healthy Doom pigs, the expression of *MYD88* is significantly lower than that in healthy control pigs. Based on the summary of Least Squares (LS) Means in the ANOVA test, it appears that there is a significant difference in the expression level of *MYD88* between ASFV infected, survived, and healthy control pigs, compared to healthy Doom pigs (Fig. 6 A1). The expression of *MYD88* in ASFV-infected pigs was found to be significantly higher than that in healthy control pigs. Additionally, a significant difference was observed in the expression level of *MYD88* between ASFV infected and survived pigs. This suggests that *MYD88* may be involved in the pathogenesis of ASFV and that differences in the expression level of this gene may be associated with differences in susceptibility to the disease. However, no significant difference was observed between healthy control and ASFV-infected pigs in *MYD88* expression, which suggests that this gene may not be involved in the initial response to ASFV infection.

**Fig. 6:**
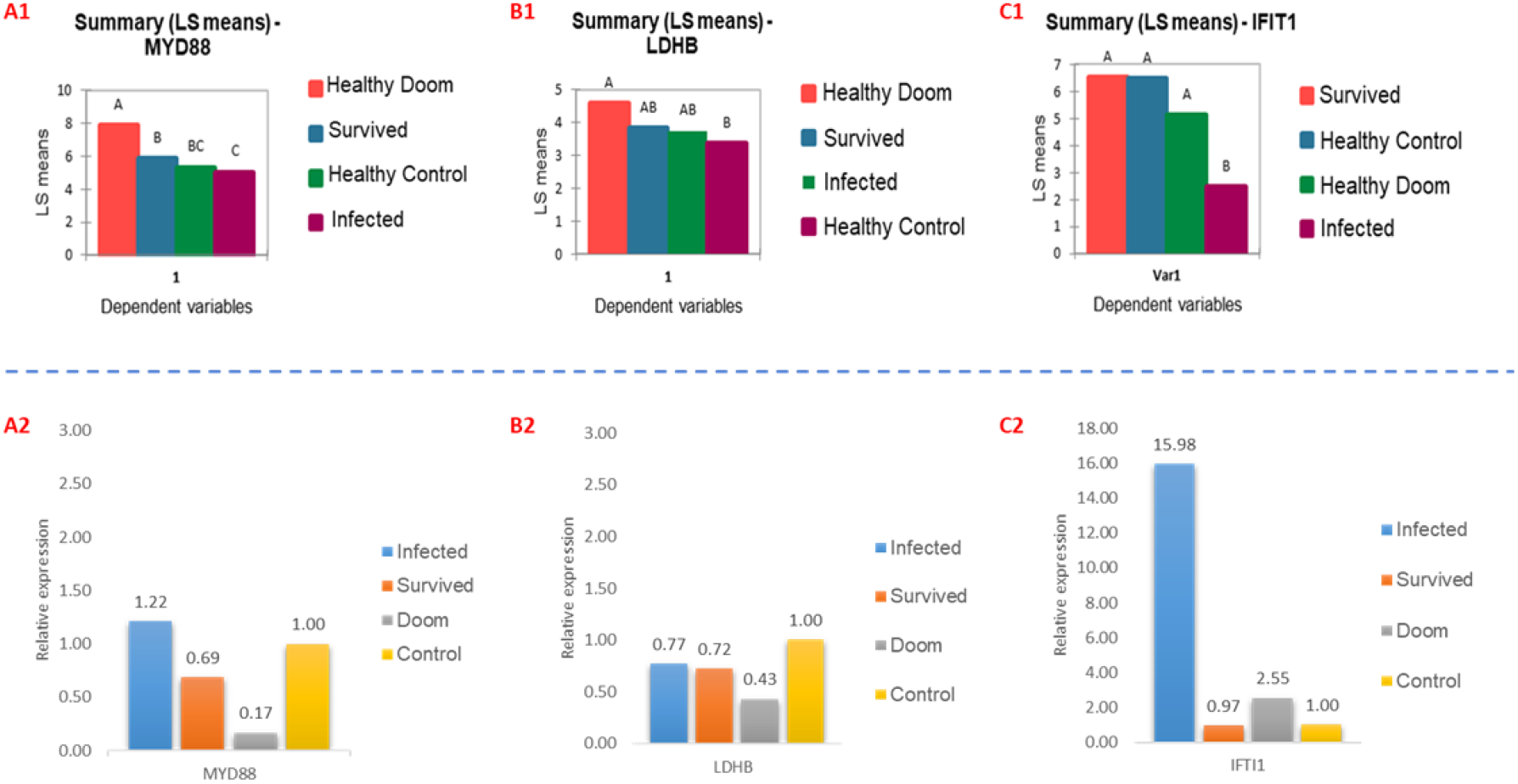
Graphical representation of the results of qPCR expression analysis of three genes, *MYD88, LDBH*, and *IFIT1*, in blood samples from different groups of pigs: healthy doom, healthy control, ASFV infected, and survived. The data is presented as the summary of least squares means in the ANOVA test of normalized ΔCT values (A1, B1, and C1) and as relative fold change values (A2, B2, and C2) for each gene in the respective experimental groups. These results provide insights into the differential gene expression patterns associated with ASFV infection and survival in pigs. Specifically, the findings suggest that *MYD88, LDBH*, and *IFIT1* might be involved in the host immune response to ASFV infection.

In the ANOVA test (summary of LS Means), *LDHB* expression levels in healthy Doom pigs were significantly different from those in healthy control pigs. However, there was no significant difference in *LDHB* expression levels between ASFV-infected and survived pigs with healthy control or healthy Doom pigs (Fig. 6 B1). Based on the qPCR analysis, *LDHB* expression was found to be downregulated in ASFV-infected pigs, with a relative fold change of 0.77. Survived pigs also exhibited reduced *LDHB* expression, with a relative fold change of 0.72, while healthy Doom pigs showed the lowest expression, with a relative fold change of 0.43 compared to healthy control (which was set as 1 fold). These findings suggest that *LDHB* may be involved in the host response to ASFV infection, with its expression being downregulated in both infected and survived pigs (Fig. 6 B2).

In ANOVA test (summary of LS Means), revealed that the expression of *IFIT1* was highly significant in ASFV-infected pigs compared to healthy control, healthy Doom, and ASFV-survived pigs (Fig. 6 C1). However, there was no significant difference observed in *IFIT1* expression among healthy control, healthy Doom, and ASFV-survived pigs. There may be a basal level of *IFIT1* expression that is not affected by ASFV infection or by differences in genetic background or other factors among the healthy and ASFV-survived pigs. The relative fold change of *IFIT1* expression (Fig. 6 C2) was significantly upregulated in ASFV-infected pigs, with a fold change of 15.98. However, survived pigs showed a nearly unchanged expression level of *IFIT1*, with a fold change of 0.97, while healthy Doom pigs exhibited a moderate upregulation, with a fold change of 2.55 compared to healthy control (which was set as 1 fold). These results suggest that *IFIT1* may play a crucial role in the host response to ASFV infection, with its expression being significantly upregulated upon infection. However, the lack of upregulation of *IFIT1* expression in survived pigs suggests that the downregulation of this gene may contribute to the successful host defence against ASFV infection.

## 4. Discussion

The study was conducted to investigate the potential role of specific genes in the innate immune response to ASFV infection in pigs. The selection of the Doom pig breed as a potential resistant breed was based on previous studies that had shown that the African wild pig, which has a composite haplotype relationship with the Asian wild pig, was a natural reservoir for ASFV and had developed resistance to the virus (4). Doom pigs were chosen because they are believed to be the closest domesticated pig breed to the wild pig (Data not published). Moreover, the use of Doom pigs as a control group in ASFV infection studies is a reasonable approach since there is no previous report on ASFV infection in this breed. This strategy allows for the identification of genetic and/or immunological factors that may contribute to ASFV resistance in other pig breeds.

The present study also conducted an in-depth analysis of *B646L* (p72) gene sequences to identify any potential genetic diversity among circulating African Swine Fever Virus (ASFV) strains in Northeast India and other parts of the world. All samples collected from e different locations of Northeast India during 2021-2023 showed typical clinical symptoms of ASFV infection as well as confirmed by PCR test and subsequently entire *B646L* (p72) gene was amplified, purified, and sequenced. Web-based BLAST and MEGA XI software were used for sequence analysis. Analysis of the *B646L* (p72) gene showed that all nine ASFV isolates from countries like China, Vietnam, Nigeria, Estonia, Armenia, Germany and Korea belonged to genotype II and were 100% identical to sequences of the present study as well as previously identified ASFV strains in India (9). Analysis of the *B646L* (p72) did not indicate any change in the nucleotide and amino acid sequences among these strains in 2 years of research and no novel variant was found. The results of this study revealed that *B646L* (p72) gene showed 100% similarity with ASFV isolates detected previously in other countries. Overall the results of the present study and a previous study (10-12) showed high stability and no genetic diversity in the *B646L* (p72) gene in the ASFV genome. One study conducted *B646L* (p72) gene in China also shows that ASFV genotype II is predominant among the 24 genotypes (13). The main goal of the present study was to identify whether tolerances to ASFV in Doom Pig was due to variation in DNA sequences *B646L* (p72) gene or difference in the host receptor site. It will be interesting to reveal any variation in the host receptor site in future studies. However, the study indicated that there might be an epitope of the protein differences in the *B646L* (p72) gene that causes variation in infection (2). This study on *B646L* (p72) gene also confirmed that there is no co-circulation of ASFV genotypes in India, which may further over cum complications to arise in ASF control measures. But this study was conducted only on three samples, therefore, it is recommended that more sample need to be sequenced for the *B646L* (p72) gene to strengthen the present finding that in India only Genotype II of ASFV is in circulation without any variation in DNA sequences as well as tolerances to ASFV in certain indigenous pig like Doom is due to not because of sequence variation *B646L* (p72) gene but of the variation of the host receptor site or there immunologically important gene.

In this connection four groups of animals (pigs) such as I. Healthy exotic breed, II. Healthy indigenous breed, III. Pigs infected with ASF and IV. Pigs that survived ASF were taken for the present study. ASFV-infected, recovered and healthy groups were confirmed by the amplification of the ASFV *B646L* (p72) gene using PCR. The survival and resistance of ASFV confirmation were done by detection of the antibody in ELISA. The presence of antibodies in pigs infected with ASFV is influenced by the virulence of the viral strain (14). Highly virulent strains often result in pig mortality before specific antibodies can be produced (15). The timing of antibody production following ASFV infection varies. Antibodies can appear 7-10 days after infection and may persist for months or even years (14, 15). Studies by Petrov et al. (2018) and Walczak et al. (2021) provide further evidence of the early development and long-term maintenance of ASFV-specific antibodies (16, 17). The current study corroborates these findings, as none of the infected pigs had antibodies during the early symptomatic phase, but two pigs developed antibodies following severe infection, aligning with the delayed antibody response observed in highly virulent cases. Simultaneously, the biomarker/ indicator genes such as *MYD88, LDHB* and *IFIT1* were observed in the DNA of blood samples of pigs in all 4 groups. The expressions of these three genes were also observed using RNA from the blood of all 4 groups of animals using Qualitative RT PCR as well as quantitative RT-PCR.

*MYD88* is involved in signalling within immune cells (18). It acts as an adapter that connects proteins that receive signals from outside the cell to the proteins that relay signals inside the cell. It is important for an early immune response to foreign invaders (antigen) (19). So, the expression of *MYD88* in the ASFV-infected group was found to be higher than in healthy exotic animals (control) but the expression level reduced in recovered and healthy Doom breed of pigs. *MYD88* may be involved in the modulation of the immune response to ASFV. *MYD88* expression was upregulated in porcine alveolar macrophages after infection with a highly virulent ASFV isolate (20). *MYD88* may have a negative effect on the antiviral type I interferon response by interfering with other signalling pathways. Therefore, targeting *MYD88* with small molecule inhibitors may have therapeutic potential for modulating host immunity against inflammatory diseases and antiviral therapy.

Secondly, *LDHB* is a terminal metabolic enzyme in the glycolysis pathway that has been widely studied in cancer cells but its role in viral infection is relatively unknown (21). A study has shown that inhibition of *LDHB* inhibited apoptosis, providing an environment conducive to persistent viral infection. It was also demonstrated that *LDHB* inhibition promoted CSFV growth through mitophagy, whereas its overexpression decreased CSFV replication. In the present study, the expression of the *LDHB* gene is lower in ASFV-infected and recovered and healthy Doom breed of pigs than the healthy control group. But Feng *et al*. have confirmed that *LDHB* has the ability to hinder the replication of ASFV (21).

Innate immunity is the first line of defence against invading pathogens. Interferon is a protein produced by a variety of cells during the process of virus infection (22). IFNs significantly trigger the production of a variety of IFN-induced genes such as IFIT family genes (22) which are involved in regulating innate immune responses. They are important targets with potent antiviral activities. They could restrict various viruses, stimulate apoptosis and regulate immune responses (23). IFTI family proteins are predominantly induced by Type I and Type III interferon and are regulated by pattern recognition and the JAK-STAT signalling pathway (23). IFIT family proteins are involved in many processes in response to viral infection; however, viruses can escape the antiviral functions of the IFIT family by suppressing the IFIT family gene expression (23). *IFIT1* is an IFNB-dependent gene, which shows significant expression in ASFV-infected and healthy Doom breeds of pigs than a healthy pig, while expressed lower than in healthy pigs in survived ASF group. In the infected group of ASFV might be induced the production of INFB-dependent genes *IFIT1*. But, increased viral load during time ASFV blocks IFNB mRNA synthesis (24). Yang *et al*. reported that *IFIT1* was significantly upregulated in ASFV-infected porcine alveolar macrophages (PAM). Activation of the *IFIT1* indicates that PAMs are in an antiviral state, which is varied with activation of the immune-related pathways (25). *IFIT1* genes for viral defence were negatively correlated with the viral load during infection which suggests that antiviral genes continued to be suppressed with high levels of viral loads in infected cells (26). ASFV E120R protein inhibits INFB production by interacting with IRF3 to block its activation to the recruitment for TANK binding kinase (TBK1) (27). Garcia-Belmonte *et al*. reported *IFIT1* expression during virulent ASFV Armenia/07 infection does not activate but attenuated NH/P68 infection expresses *IFIT1* significantly. The virulent Armenia/07 strain is able to efficiently block INFB mRNA synthesis by inhibition of IRF3 activation, STING phosphorylation and production in infected macrophages, whereas NH/P68 does not (28). Out of these 3 genes *IFIT1* was expressed higher in the Doom pig than control. In the case of the Doom breed of pig, it is genetically close enough to wild pigs and has also a scavenging nature like wild pigs. A recent study also shows that Kenya domestic pigs which are pure indigenous pigs of Kenya also showed the same kind of resistance to ASFV as its wild counterpart. The detection of ASFV was also seen to correlate with pig genotype, with a higher proportion of pigs with low local African ancestry testing positive for the virus. A comparative study on selection signatures between the wild African and local domestic African pigs indicates the absence of broad patterns of selection that are similar between the domestic and wild pigs. These indigenous pigs particularly found in Africa and Asia are the natural habitat of pigs’ domestication and origination and they are more closely related to their wild ancestors than the modern European and American breeds. Few studies also show that African pigs have a composite ancestral relation with haplotypes from Asian counterparts. In the Asian context, particularly, in the Northeastern region of India which is one of the main hotspots for pig origination and domestication has some prized indigenous pig breeds. Among total registered pig breeds Doom pig from North East India is present in breeding tracks having latitude and longitude of 26.0°to 26.6°N and 89.3° to 90.4°E with an approximate area of distribution of about 3,000 km^2^ not only the closest progenitor of Indian wild pig (unpublished data) but also very much adaptable to harsh environmental conditions and thrives with very low to limited nutritional input and is capable of surviving in a migratory scavenging system (29). Because of having typical hardiness and coping power with common pig diseases and closeness with wild counterparts this pig breed is the typical candidate of pig breed to tolerant to ASFV infection. So, the present study investigates the natural resistance to African swine fever in Doom pigs from an endemic area in North East India to support the proposition that Doom pigs can co-exist with virulent ASF viruses.

## 5. Conclusion

ASFV is a serious threat to the piggery sector in India as a whole and particularly the northeastern part of the country where the majority of the pig population exists. Due to a lack of vaccines, and biosecurity measurements, other scientific efforts are being made to control the spread of the disease. However, in the cross-country movement of pork and pig, ASFV becomes endemic in the Country. The development of a vaccine and identifying or producing the tolerant or resistant pig to ASFV will be a great help in controlling the disease as well as repopulating the lost population. Maybe in the long run, genetically modified pigs with resistance power to ASFV will be developed. But acceptance among consumers of GM pigs is also questionable. Since few past and current research also showed that some of the wild pig members carry certain transcriptomic factors that show resistance to ASFV, it was an urgent need to study ASFV resistance patterns in indigenous Doom pigs of the region. Overall a comprehensive approach at the genomic/ transcriptomic level is required to explore the possible genetic determinants underlying tolerance to infection by ASFV genotypes and identify genetically mediated ASFV tolerance factors/genes in the indigenous population of Pigs. The results of this study revealed that the *B646L* (p72) gene showed 100% similarity with ASFV isolates detected previously in other countries as well as confirmed that there is no co-circulation of different genotypes of ASFV in India. Present findings also suggested that *MYD88, LDHB* and *IFIT1* are involved in the pathogenesis of ASFV and that their expression pattern will be a useful biomarker for distinguishing between different stages of the disease. However, further research is needed to confirm these findings and to understand the underlying mechanisms involved in the differential expression of *MYD88, LDHB* and *IFIT1* in ASFV infected, survived, and healthy control pigs compared to healthy Doom pigs.

## Author Contributions

PJD conceived and designed the research. PJD, JS, GSS and SRP conducted the wet lab work. PJD and JS analyzed the data. PJD, JS, SRP RD and SK helped in manuscript drafting and editing. PJD, SRP, SR, SB and VKG supervised and provided the resources. All authors revised and approved the final version of the manuscript.

## Conflict of Interest

The authors declare that they have no conflict of interest.

## Data Availability Statement

The data that support this study will be shared upon reasonable request to the corresponding author.

## Funding

The study was supported by Indian Council of Agricultural Research-National Research Centre on Pig Institutional Project Grant 2022-2023.

## Acknowledgements

The authors are thankful to the Director, Indian Council of Agricultural Research-National Research Centre on Pig for granting permission to carry out this project.

